# Multiplexed quantum sensing reveals coordinated thermomagnetic regulation of mitochondria

**DOI:** 10.1101/2025.07.30.666664

**Authors:** Md Shakil Bin Kashem, Stella Varnum, Olivia Lazorik, Rocky Giwa, Shiva Iyer, Changyu Yao, David W. Piston, Chong Zu, Jonathan R. Brestoff, Shankar Mukherji

## Abstract

Mitochondria are multifunctional organelles that convert the potential energy stored in nutrients and intermediary metabolites into both heat and an electrochemical proton-motive force. However, how these outputs are synchronized in cells remains an enduring question. In this work, leveraging multiplexed nanodiamond quantum sensors to monitor both changes in temperature and magnetic field fluctuations in single primary cells obtained from diverse tissues in adult mice, we identified thermomagnetic correlation profiles uncovering a regulatory feedback loop in which the cell draws upon available intracellular iron to maintain the mitochondrial electrochemical gradient. These profiles reverse in cells derived from a mouse model of Leigh syndrome and raise the intriguing possibility that primary mitochondrial diseases can be understood as disorders of thermomagnetic homeostasis.

## Introduction

Mitochondria are essential for cellular function, playing a central role in energy production, maintaining redox balance, and regulating iron homeostasis(*1–4*). The energy production process of oxidative phosphorylation is driven by an electrochemical gradient generated across the inner mitochondrial membrane. Proton transport across the membrane is coupled to electron transfer from embedded membrane complexes I-V, ultimately generating adenosine triphosphate (ATP), the primary cellular energy currency (*1*). However, uncoupling the dissipation of the proton gradient from ATP synthesis results in the generation of substantial levels of heat. Brown adipocytes employ uncoupling by expressing Uncoupling protein 1 (UCP1) to generate heat in a processs called nonshivering or adaptive thermogenesis (*5–8*). A complete account of mitochondrial proton pumping regulation is of fundamental importance to both cell biology and clinical medicine, especially for metabolic diseases characterized by impaired mitochondrial metabolism.

The electron transport chain (ETC) that is used to generate the mitochondrial proton gradient is comprised of 5 protein complexes, some of which contain iron-sulfur (Fe-S) clusters that are essential for oxidative phosphorylation. As a result, mitochondria are a primary site for iron utilization within the cell, where imported iron is critical for both heme synthesis and Fe-S cluster biogenesis (*9*). Mitochondrial dysfunction, including increased generation of reactive oxygen species (ROS) and reduced oxidative phosphorylation, is implicated in the pathophysiology of metabolic disorders, including primary mitochondrial diseases such as Leigh syndrome and secondary mitochondrial diseaes such as obesity, type 2 diabetes, and heart failure (*1*). Of particular interest is the interplay among mitochondrial thermogenesis, Fe-S cluster biogenesis (*10*), ROS generation via the ETC, and iron homeostasis (*4, 11, 12*). A deeper understanding of how cellular temperature relates to iron availability requires a sensor capable of simultaneously monitoring both.

Conventional techniques for the sensing of intracellular temperature, iron, and ROS levels, namely fluorescent dyes, are limited by photobleaching and prone to artefactual signal changes (*13– 15*). Further pitfalls include the difficulty of calibration, complicated by temperature-independent factors in the complex cellular environment and incomplete dye hydrolysis (*14, 15*). In addition, by reacting reversibly with ROS, these dyes reveal the history of the sample rather than its current state (*16*). Therefore, there is a need to explore alternative sensing methods for improved sensitivity and subcellular resolution (*17*).

Nitrogen-vacancy (NV) centers are atomic-scale defects in diamond whose spin levels are sensitive to local variations in temperature and magnetic noise, making them powerful tools for nanoscale sensing in biological environments (*18–37*). Their biocompatibility and dual sensitivity enable multiplexed measurements of thermal and magnetic signals, with a few recent studies demonstrating such capabilities in immortalized cell lines (*25, 38*). However, to our knowledge, no prior study has applied multiplex quantum sensors in primary cells to uncover biologically meaningful, co-regulated metabolic and iron dynamics at the single-cell level.

In this work, we leverage the multiplex sensing capability of nanodiamond quantum sensors containing NV centers to measure correlated changes in temperature and magnetic noise within different primary cell types extracted from adult mice, uncovering regulatory mechanisms relating mitochondrial proton pumping and subcellular iron utilization.

### Multiplexed nanodiamond quantum sensor of temperature and magnetism

Primary immune cells were harvested from the peritoneal cavity of mice and incubated with 70 nm nanodiamonds (Fig. 1a) (*39*). Following cellular uptake, residual nanoparticles were removed by thorough washing (see Supplementary Materials for detailed protocols on cell isolation and nanodiamond preparation). Fluorescence from intracellular nanodiamonds was recorded with a custom-built confocal microscope (Fig. S1, S2, (*40*)). A representative bright-field image of a single cell is shown in Figure 1b, with the corresponding confocal raster scan of nanodiamond fluorescence displayed in Figure 1c.

**Figure 1:**
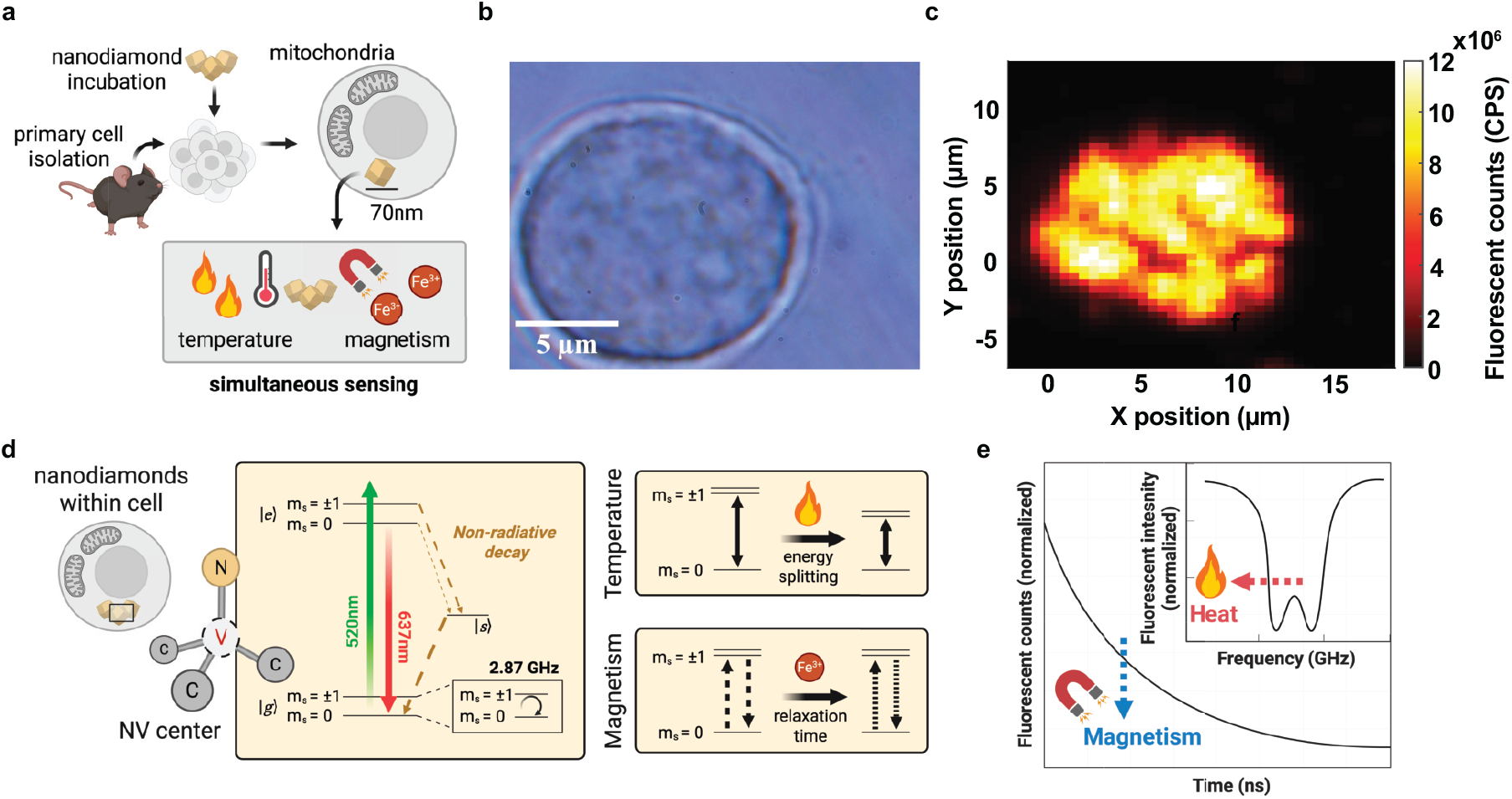
Nanodiamond uptake by mouse primary cells enables dual sensing of magnetism and temperature. (a) Experimental design: mouse primary cells were isolated and incubated with 70 nm nanodiamonds to enable dual temperature and magnetic sensing. (b) Brightfield image showing live primary cells with internalized nanodiamonds. (c) Confocal fluorescence raster scan verifying intracellular nanodiamond localization. (d) Crystal lattice structure and electronic energy level diagram of a single nitrogen-vacancy (NV) center. Raising intracellular temperature lowers the ground state energy splitting between *m*_*s*_ = 0 and *m*_*s*_ = ±1 levels, while enhanced magnetic fluctuations lead to faster spin relaxation between the levels and shorten the *T*_1_. (e) Schematic of the measured ODMR spectrum (inset) and *T*_1_ relaxation curve of nanodiamond NV centers for correlating intracellular temperature and magnetism.

Each nanodiamond NV center consists of a substitutional nitrogen impurity adjacent to a vacancy, replacing two intrinsic carbon atoms within the diamond lattice. The electronic ground state of the NV center exhibits a spin-1 degree of freedom (Fig. 1d). In the absence of an external field, |*m*_*s*_ = ±1⟩ spin levels are near degenerate and separated from |*m*_*s*_ = 0⟩ by the zero-field splitting *D* = 2.87 GHz. This spin transition energy *D* can be probed using optically detected magnetic resonance (ODMR) spectroscopy: by sweeping the frequency of the applied microwave drive while monitoring the fluorescent signal, we expect a decrease in fluorescence when the microwave frequency is resonant with *D* (Fig. 1e) (*19, 41–44*). When the local temperature of the nanodiamond changes, *D* exhibits a well-characterized, linear, temperature-dependent shift *dD* (*T*)/*dT* ≈ −70 kHz/^*o*^C, enabling the measurement of subcellular temperature (Fig. S3, (*40*)).

Concurrently, fluctuating magnetic noise from nearby Fe^3+^ can induce transitions between spin states and shorten the NV lifetime, *T*_1_ (Fig. S4, (*40*)). By tracking both *D* and *T*_1_, we can therefore correlate intracellular temperature fluctuations with changes in ferric iron concentration (Fig. 1e).

### Thermogenic responses in mouse peritoneal exudate cells

We started by measuring the intracellular temperatures in individual mouse peritoneal exudate cells (PECS) collected by lavaging the abdominal cavity of euthanized mice (Fig. 2a) (*39*). After incubating the cells with nanodiamonds, we either left them untreated (CTRL) or exposed them to 2 *µ*M carbonyl cyanide-p-trifluoromethoxyphenylhydrazone (FCCP) immediately before sensing. FCCP is a mitochondrial protonophore that collapses the inner membrane electrochemical gradient, releasing the stored energy as heat rather than coupling it to ATP synthesis; it is therefore expected to raise cellular temperature (Fig. 2b) (*45–48*).

**Figure 2:**
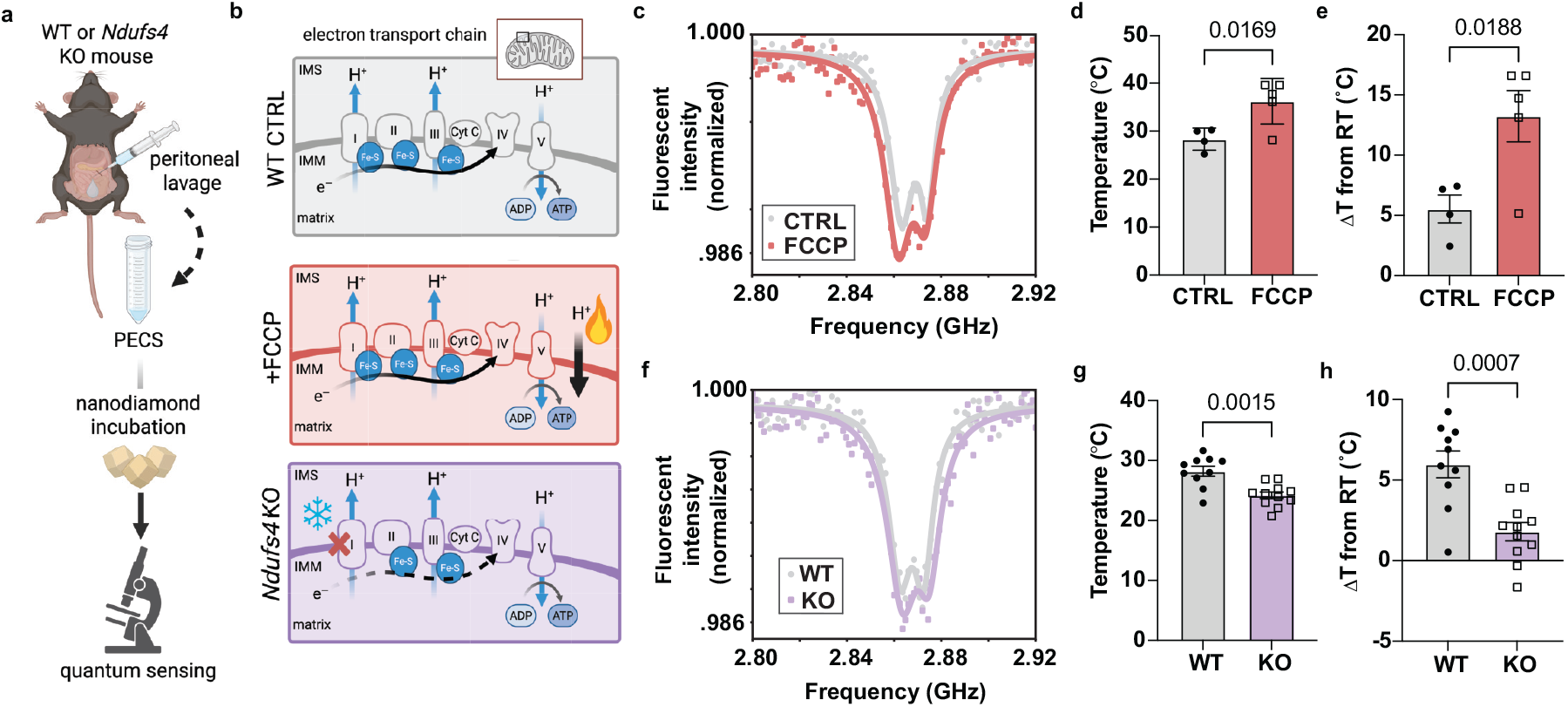
Quantum sensing of mouse peritoneal exudate cells (PECS) reveals temperature fluctuations in response to both pharmacologic and genetic manipulation of proton pumping. **(a)** Isolation of PECS from wild-type (WT) or *Ndufs4*^−/−^ (KO) mice. PECS were collected by peritoneal lavage, incubated with nanodiamonds, and then either treated with FCCP or left untreated (CTRL) prior to quantum sensing. (b) Effect of FCCP treatment and *Ndufs4* deletion on mitochondrial electron transport chain function. FCCP uncouples proton transport from ATP synthesis, dissipating the gradient and generating heat. In contrast, *Ndufs4* deletion impairs complex I assembly, reducing proton gradient formation and thermogenesis. (c) Representative ODMR spectra for CTRL and FCCP-treated PECS. (d) Mean intracellular temperature from the ODMR spectra (*n* = 4 CTRL, *n* = 5 FCCP). (e) Change in intracellular temperature relative to room temperature baseline (Δ*T*). (f) Representative ODMR spectra for WT and KO PECS. (g) Mean intracellular temperature from the ODMR spectra (*n* = 10 WT, *n* = 11 KO). (h) Δ*T* for WT and KO PECS relative to room temperature. Individual data points are shown alongside error bars representing group means ±SEM. Statistical significance in (d), (e), (g), and (h) was performed using Student’s t-test.

Figure 2c shows representative ODMR spectra from a CTRL cell and an FCCP-treated cell. Indeed, in the FCCP condition, the zero-field splitting *D(T*) was shifted to lower microwave frequencies, indicating an increase in intracellular temperature. Averaging across multiple cells, we found that FCCP treatment raises the mean cellular temperature to 36.2 ± 2.1 °C, higher than the 28.3 ± 1.2 °C measured in CTRL cells (Fig. 2d). To more clearly illustrate the thermogenic output relative to the environmental baseline, we computed the change in intracellular temperature (Δ*T*) from the room temperature recorded at the time of measurement (≈ 22 °C). As shown in Figure 2e, FCCP-treated cells exhibited a significantly larger Δ*T* than CTRL, confirming that the observed temperature rise reflects active cellular heat generation. Our result highlights the utility of NV thermometry for capturing metabolic heat dynamics within single primary cell resolution.

FCCP is a well-established stimulus for dissipating an established mitochondrial proton gradient to generate heat. Therefore, we sought a complementary approach of genetically disrupting proton pumping to prevent normal formation of the proton gradient using *Ndufs4*^−/−^ (KO) mice. NDUFS4 is a subunit of mitochondrial complex I, and loss of function of this gene impairs complex I-dependent electron transport and proton pumping, leading to the development of a fatal primary mitochondrial disease called Leigh syndrome (Fig. 2b) (*49, 50*). To this end, we examined PECS isolated from *Ndufs4*^−/−^ (KO) mice to assess how chronic mitochondrial dysfunction impacts intracellular temperature. ODMR spectra from KO cells displayed an upshift in resonance frequency, indicating a lower intracellular temperature than wild-type (WT) cells (Fig. 2f). Averaged over multiple cells, KO temperature, 24.2 ± 0.6 °C, was much lower than WT (Fig. 2g). We further calculated Δ*T* for WT and KO PECS to quantify thermogenic output relative to the baseline room temperature. As shown in Figure 2h, WT exhibited significantly higher Δ*T* than KO counterparts (*p* = 0.0007), indicating impaired mitochondrial heat production in the context of *Ndufs4* deficiency. This temperature deficit aligns with the known role of complex I in maintaining oxidative phosphorylation and underscores NV-based thermometry as a sensitive reporter of mitochondrial function.

### Iron-driven coordination of thermogenesis in splenic immune cells

Having established mitochondrial activity–dependent temperature differences in both pharmacologically and genetically perturbed PECS, we next leveraged the multiplex sensing capabilities of nanodiamonds to simultaneously measure intracellular temperature and ferric iron levels, aiming to uncover mechanistic links between mitochondrial proton pumping and heat generation. We chose to work with cells from the mouse spleen, focusing on a magnetic subset particularly enriched in ferritin and ferric iron (Fe^3+^) (Fig. 3a). In this subpopulation of iron-rich cells, the iron level is sufficient to allow their purification via magnetic columns (see Supplementary Materials for full details of splenocyte isolation and nanodiamond loading protocols). We referred to these as magnetic cells and compared them to splenic cells that were not enriched via magnetic column, thereby termed non-magnetic (Fig. 3a).

**Figure 3:**
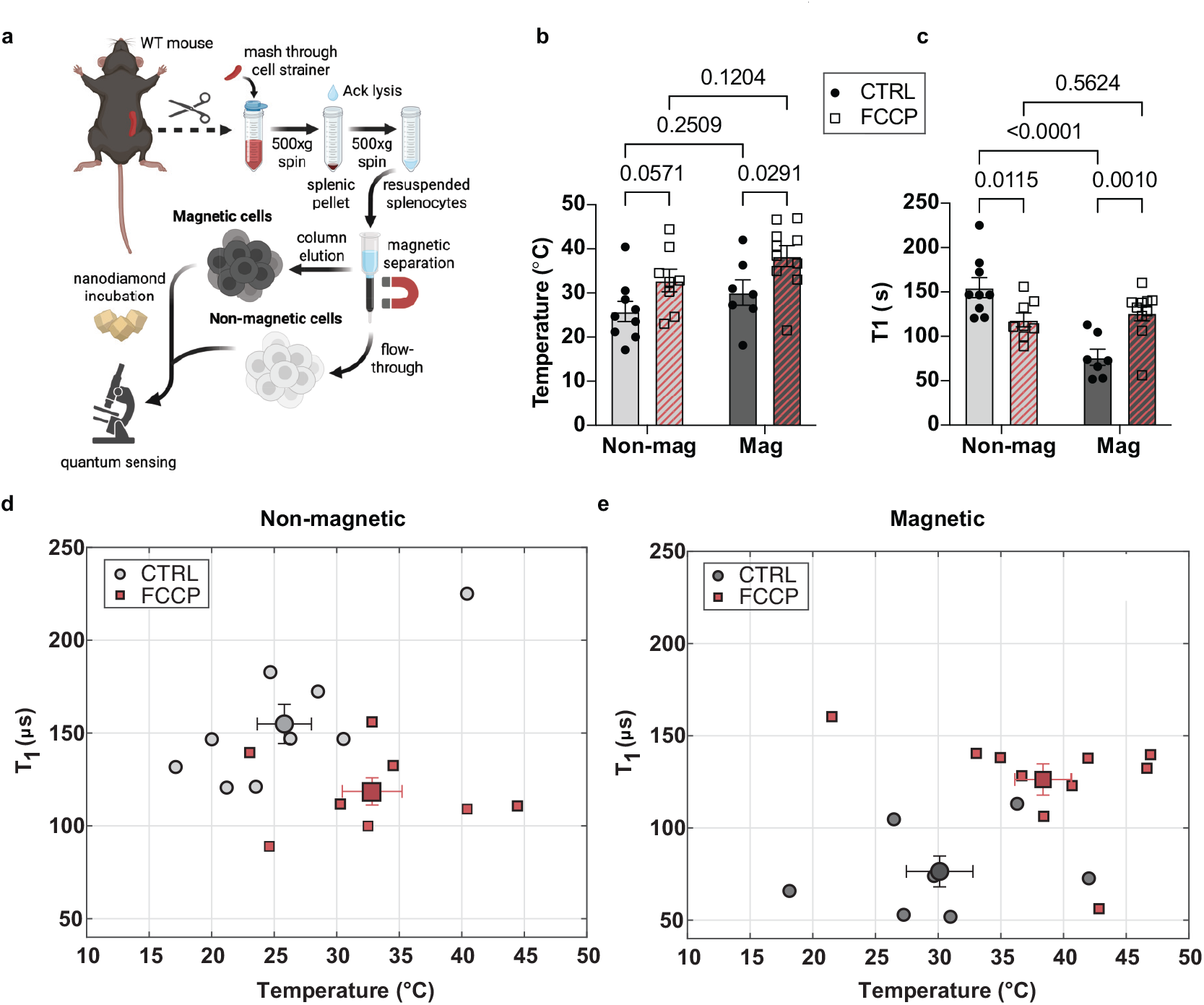
Mitochondrial uncoupling modulates thermomagnetic correlation profile in WT splenic cells. (a) Isolation of splenic cells from wild-type (WT) mice, followed by magnetic column separation of iron-rich (magnetic) and unretained (non-magnetic) populations, treatment with FCCP, and measurement with NV-based quantum sensing. (b) Mean intracellular temperatures of WT magnetic (mag) (*n* = 7 CTRL, *n* = 10 FCCP) and non-magnetic (non-mag) (*n* = 9 CTRL, *n* = 8 FCCP) splenic cells, with and without FCCP treatment. (c) Corresponding mean NV spin relaxation times (*T*_1_) for the same groups. (d) Scatter plot correlating intracellular temperature with *T*_1_ for individual non-magnetic and (e) magnetic WT splenocytes. Individual data points are shown alongside error bars representing group means ±SEM. Statistical analysis: two-way ANOVA with repeated measures.

In each of these populations of splenocytes, we deployed multiplexed nanodiamond quantum sensing to acquire ODMR spectra, followed by *T*_1_ relaxometry measurements to assess the combined thermal and magnetic response. For thermometry, magnetic cells exhibited an average temperature of 30.1 ± 2.9 °C while non-magnetic cells were slightly cooler at 25.8 ± 2.3 °C. However, this temperature difference falls within the experimental uncertainty (Fig. 3b). In contrast, *T*_1_ relaxometry revealed a pronounced difference: the magnetic cells showed a significantly shorter average *T*_1_ of 76.4 ± 9.0 *µ*s compared to 154.9 ± 11.2 *µ*s in non-magnetic cells (*p* < 0.0001), (Fig. 3c), consistent with the elevated paramagnetic spin noise in iron-rich cells.

Upon FCCP treatment, magnetic cells showed a significant increase in temperature to 38.4 ± 2.4 °C (Fig. 3b,e), accompanied by a *T*_1_ increase to 126.3 ± 8.9 *µ*s (*p* = 0.0010) (Fig. 3c),reflecting the behavior of single cells (Fig. 3e). Interestingly, in non-magnetic splenic cells, we observed the opposite trend: FCCP treatment resulted in a modest reduction in *T*_1_ to 118.6 ± 7.9 *µ*s (*p* = 0.0115)(Fig. 3c,d), despite increased temperature (32.8 ± 2.6 °C) (Fig. 3b), also reflected in the behavior of single cells (Fig. 3d).

Given these findings, we hypothesized that an iron-mediated effect could account for the observed anti-correlated (Fig. 3d) and correlated (Fig. 3e) responses in *T*_1_ and temperatures to FCCP. Our model rests on the idea that ETC activity adjusts to maintain the mitochondrial proton gradient. From this assumption, we derive two possibilities:

1. Correlated *T*_1_ and temperature: By dissipating the mitochondrial membrane potential, FCCP may trigger compensatory mechanisms that increase Fe-S biogenesis by drawing down from the pool of ferric iron in the cell to restore the proton gradient.
2. Anti-correlated *T*_1_ and temperature: FCCP-induced uncoupling might lead to increased electron flow through the ETC to restore the gradient. Heightened ETC activity could elevate mitochondrial ROS levels. Elevated ROS can additionally lead to oxidative degradation of Fe-S proteins via the Fenton reaction, which cannot be counteracted by the available pool of subcellular iron to support increased Fe-S biogenesis (*1, 4, 51*).

A key prediction of this model is that if Fe^3+^ levels in non-magnetic splenic cells increase, for example by a shift in the equilibrium of the subcellular iron pool from ferrous iron (Fe^2+^) or Fe-S clusters to ferric iron, it is possible that the *T*_1_ versus temperature correlation will reverse signs from negative to positive. Inhibiting complex I, a key ETC protein composed of Fe-S, has been studied and found to decrease Fe-S cluster biogenesis (*52*). In addition, depletion of complex I on the organismal level, as seen with Ndufs4 KO mice, leads to perturbations in iron utilization indicative of iron overload in these mice, whereas iron restriction leads to reduced disease severity (*53*). Should the non-magnetic splenocytes from *Ndufs4*^−/−^ knockout (KO) mice exhibit decreased rather than increased *T*_1_ coincident with a decreased temperature, this could indicate that the iron equilibrium in these cells have shifted away from Fe-S toward the stored form of ferric iron. Ferric iron would then be available to upregulate Fe-S formation upon treatment with FCCP and cause a type switch of the non-magnetic cells from having an anti-correlated to correlated *T*_1_ versus temperature change. To test this prediction, we performed the same measurements in splenocytes isolated from *Ndufs4*^−/−^ KO mice (Fig. 4a). Consistent with our results from peritoneal cavity cells, both magnetic and non-magnetic KO splenocytes exhibited lower intracellular temperatures than WT controls. Untreated magnetic KO cells showed an average temperature of 27.7±2.6 °C, rising to 34.1±1.5 °C upon FCCP treatment (*p* < 0.0554) (Fig. 4b,e). Non-magnetic KO cells similarly increased in temperature from 21.1 ± 1.9 °C to 33.8 ± 1.9 °C after FCCP treatment (*p* < 0.0001) (Fig. 4b,d).

**Figure 4:**
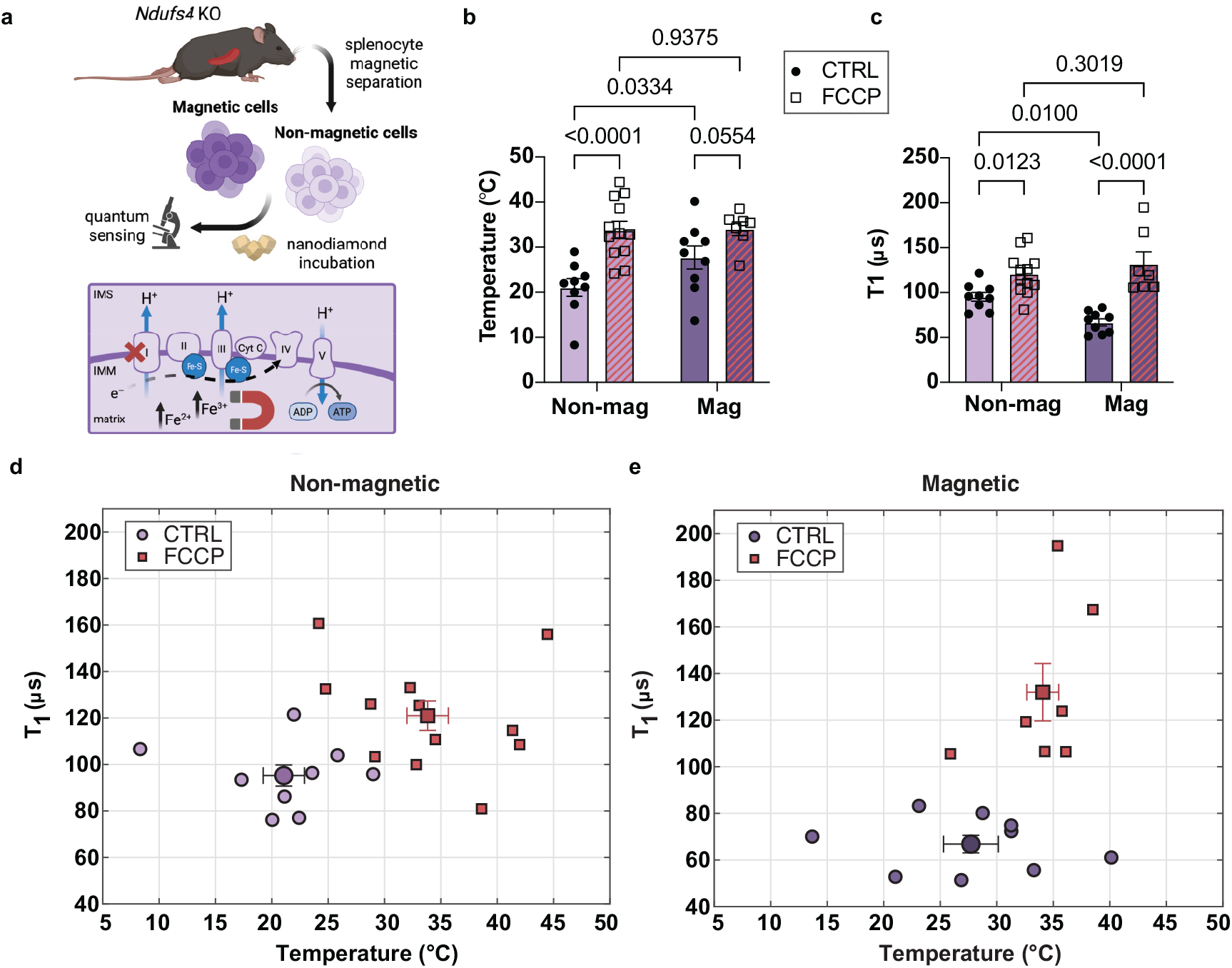
Non-magnetic splenocytes from Ndufs4 KO mice exhibit reversed thermomagnetic correlation profiles compared to WT. (a) Experimental schematic highlighting Ndufs4 knockout (KO) cell isolation for quantum sensing, as well as ETC disruption in KO cells and expected changes in thermogenesis and redox signaling. (b) Mean intracellular temperatures of KO nonmagnetic (*n* = 9 CTRL, *n* = 12 FCCP) and magnetic (*n* = 9 CTRL, *n* = 7 FCCP) splenic cells, with and without FCCP treatment. (c) Mean *T*_1_ relaxometry time showing magnetic noise levels across conditions. (d) Scatter plot correlating intracellular temperature with *T*_1_ for individual nonmagnetic and (e) magnetic KO splenocytes. Data points represent individual cells, with overlaid bars indicating mean ±SEM. Statistical comparisons were performed using two-way ANOVA with repeated measures.

*T*_1_ relaxometry revealed shorter relaxation times in magnetic KO cells (66.8 ± 4.0 *µ*s) compared to non-magnetic KO cells (95.2 ± 4.8 *µ*s) (*p* = 0.01), consistent with higher paramagnetic spin noise (Fig. 4c-e). Notably, FCCP treatment elevated *T*_1_ in magnetic KO cells to 132.0 ± 13.3 *µ*s (*p* < 0.0001), paralleling the trend observed in WT. As a crucial test of our model, non-magnetic KO cells also exhibited a significant increase in *T*_1_ upon treatment with FCCP (121.0 ± 6.6 *µ*s; *p* = 0.0123), (Fig. 4d) which contrasts with the behavior observed in WT non-magnetic cells (Fig. 3d). This result suggests that in ferritin-rich macrophages, FCCP-induced dissipation of the proton gradient may stimulate compensatory Fe-S biogenesis to restore ETC function, depleting ferritin stores via Fe^3+^ to Fe^2+^ conversion and thus reducing local magnetic noise.

## Discussion

Our work demonstrates the first multiplexed intracellular measurement of temperature and magnetic noise in primary cells using NV centers in nanodiamonds. This dual-mode quantum sensing strategy enables direct observation of the interplay between mitochondrial thermogenesis and magnetism in live primary mammalian cells. Uncovering feedback regulatory principles underlying mitochondrial function, especially with regard to the roles played by ionic cofactors such as iron, has been notoriously difficult. Multiplexing ODMR thermometry and *T*_1_ relaxometry in the same cell has allowed us to propose and test models for a relationship between iron and mitochondrial homeostasis.

Our results show that FCCP treatment leads to increased intracellular temperatures across all measured cells, consistent with prior reports of proton gradient dissipation and thermogenic heat release (*46–48, 54*). However, *T*_1_ measurements revealed divergent trends between magnetic (iron-rich) and non-magnetic cell populations. Magnetic cells exhibited increased *T*_1_ relaxation times following FCCP treatment, consistent with accelerated (Fe^3+^ → Fe^2+^) conversion to support compensatory Fe-S biogenesis under proton gradient loss. Conversely, non-magnetic cells showed a reduction in *T*_1_ values, suggesting that oxidative destabilization of Fe-S clusters and impaired mitochondrial metabolism may predominate under these conditions. Guided by our multiplexed quantum sensing, we were able to test our model of the role of available iron to support mitochondrial proton pumping homeostasis by measuring spin relaxation and temperature in cells with a destabilized complex I of the ETC. We observed the ‘type switch’ behavior expected by the model, in which the correlation between the spin relaxation time and temperature in the non-magnetic splenocytes resembled the correlation from magnetic splenocytes.

These findings highlight the unique capability of NV center-based quantum sensors to dissect cellular heterogeneity in iron metabolism and redox state with nanoscale spatial resolution. Conventional imaging or biochemical assays lack the temporal and spatial resolution to simultaneously capture intracellular biochemical and biophysical transformations at the single-cell level. Further studies can leverage additional properties enabled by multiplexed nanodiamond quantum sensing to even better characterize the physical environment of the cell. For example, establishing how subcellular temperature varies with local cytoplasmic viscosity, by exploiting the extreme photostability of nanodiamond for use as tracer particles (*55*), will enable us to understand the precise biophysical environment in which diverse biochemical reactions take place.

Looking forward, we envision extending this platform to a broader spectrum of biological systems, including primary neurons, stem cells, and cancer models, where mitochondrial dysfunction and iron dysregulation are hallmarks of disease. The distinct pattern of change in the spin relaxation versus temperature correlation upon FCCP treatment for non-magnetic splenocytes in KO versus WT cells points the way toward using these multiplexed quantum sensing data to diagnose cellular defects. Furthermore, incorporating organelle-specific nanodiamond delivery or targeted mitochondrial labeling could further refine spatial precision. Moreover, integrating NV quantum sensing with high-throughput drug screening pipelines may enable the discovery of modulators of oxidative stress, iron metabolism, and mitochondrial bioenergetics. Altogether, our work establishes a framework for multiplexed intracellular quantum measurements with the potential to transform our ability to track disease progression and therapeutic responses at the subcellular level in real time and raise the possibility that primary mitochondrial diseases can be understood as disorders of cellular thermomagentic homeostasis (*56, 57*).

## Acknowledgments

We gratefully acknowledge discussion with Leo Shmuylovich, Erik Henriksen, Kater Murch, Norman Yao, Chuanwei Zhang, and all the members of the Zu, Brestoff, and Mukherji groups.

## Data and materials availability

Source data are included with this article. Further data are available from the corresponding author upon request.

