## Supplementary Information for "Multiplexed quantum sensing reveals coordinated thermomagnetic regulation of mitochondria"

### Supplementary Materials for: Multiplexed quantum sensing reveals coordinated thermomagnetic regulation of mitochondria

### 1 Materials and Methods

#### 1.1 Experimental Setup

We use a home-built confocal microscope that integrates fluorescence excitation and collection (Fig. S1). Cells containing nanodiamonds are placed on the coverglass and sealed with epoxy, and the samples are then mounted onto a custom-designed coplanar waveguide (CPW) for microwave delivery (Fig. S2a). The center of the CPW is designed to a Ω-shape with radius 150 $\mu$m to concentrate the microwave signals. A through hole located at the center of the $Omega$-shape allows for imaging of the sample under white light (Fig. S2b). A patterned resistive heater was fabricated on the rear side of the PCB, allowing the control of sample temperature during experimental procedures (Fig. S2c). The CPW is secured onto a custom-fabricated slide holder, which is mounted on a 3-axis translation stage (Thorlabs MAX311D/M) NanoMax stage). The stage's position is controlled either manually, using the differential knobs (50 µm/rev), or remotely with piezo actuators. A 1.25 NA oil-immersion microscope objective (Olympus 100X Immersion Objective) focuses the beam to a diffraction-limited laser spot. A green laser (Cobolt 06 MLD) with $\lambda$ = 520 nm (power 0.5 mW) is used for optical excitation. Upon exiting the laser head, the laser beam reflects off of two mirrors before reflecting off a dichroic beam splitter ($\lambda$=560 nm) and onto the two mirrors of the Scanning Galvanometer (ThorLabs GVS012/M). A voltage sweep is applied to the Galvanometer to change the angle of the reflected beam and scan the laser across the objective lens's field of view. The beam then passes through a 4-f telescope with focal length $f$ = 500 mm to map the scanning beam onto the back aperture of the objective lens. The emitted fluorescence from the nanodiamonds passes through a $\lambda$ = 632.8 nm long pass filter and a $\lambda$ = 842 nm short pass filter, coupled into a single-mode fiber acting as an emission pinhole, before being detected by a single photon counting module (Excelitas Technologies). An LED white light source at the back side of the sample stage illuminates visible light onto the sample, which is then imaged onto a CMOS camera (Thorlabs CS 165) for brightfield imaging.

A signal generator (Stanford Research Systems Model SG384) produces the microwave signal, which is amplified by a microwave amplifier (Mini-Circuits ZHL-15W-422-S+), and shuttered by a switch (MiniCircuits ZASWA-2-50DRA+). It is delivered to the sample through the CPW. A data acquisition device (National Instruments USB X Series) processes the signal. A programmable multi-channel pulse generator (SpinCore PulseBlasterESR-PRO 500) with 2 ns temporal resolution gates the equipment.

#### 1.2 Mice

Wildtype C57BL/6J mice (strain #000664) and *Ndufs4+/-* heterozygotes (strain #027058) (*1*) on a C57BL/6J genetic background were obtained from Jackson Laboratories (Bar Harbor, Maine). Ndufs4-knockout (KO) mice were generated by breeding heterozygote males to heterozygote females. Male and female mice at age 8–10-weeks-old were used in similar proportions across all experiments. Mice were housed in a specific pathogen-free (SPF) barrier facility with a 12h:12h light:dark cycle, with the lights turned on at 6 am and off at 6 pm, at room temperature and target humidity range of 40-70 percent. WT mice were provided *ad libitum* access to autoclaved food and water, and KO mice were provided with hydrogel and food pellets on the cage floor to ensure their access to hydration and nutrition throughout neurodegenerative progression. The use of animals and all animal procedures were approved by the Washington University in St Louis School of Medicine Institutional Animal Care and Use Committee (IACUC) under Protocol #22-0286.

#### 1.3 Peritoneal exudate cell isolation

Mice were euthanized using isoflurane asphyxiation, and peritoneal exudate cells (PECs) were harvested by infusing 5-10 mL phosphate-buffered saline (PBS) into the peritoneal cavity and gently withdrawing the fluid into a sterile syringe. The cell suspension was centrifuged at $500 \times g$ for 5 min at 4°C, and the pellet was resuspended in cold PBS. Cells were counted using 0.1 percent trypan blue exclusion using a Countess FL and adjusted to a final concentration of $2 \times 10^5$ cells in 100 $\mu$L PBS for subsequent 70 nm nanodiamond loading at a concentration of 40 $\mu$g/mL in PBS for 30 minutes at 37 °C. Following the incubation, cells were pelleted by centrifugation at $500 \times g$ for 5 min, and the cell-free supernatant was discarded. Cells were resuspended in 200 $\mu$L of PBS and pelleted by centrifugation to remove excess nanodiamond in the media. Finally, the cells were resuspended in 100 $\mu$L of MACS buffer (PBS + 0.5 % bovine serum albumin + 2mM EDTA). For FCCP-treated cells, FCCP was added to a concentration of 2 $\mu$L immediately prior to coverglass preparation. We prepared the sample by creating a chamber of 80 $\mu$L of cells in media sealed with epoxy between two pieces of coverglass.

#### 1.4 Splenocyte isolation and magnetic sorting

Spleens were harvested from euthanized mice and were gently mashed through a 100 $\mu$m cell strainer using the plunger of a sterile syringe and washed with PBS. Cells were centrifuged at $500 \times g$ for 5 min at 4°C, and the supernatants were aspirated. Red blood cells were lysed with 3 mL of ammonium chloride potassium (ACK) lysis buffer (Lonza) for 5 min at room temperature, and the reaction was quenched by adding 10 mL PBS and gently mixing. The cells were pelleted by centrifugation at 500 x *g* for 5 min at 4°C, resuspended in 10 mL PBS to wash off residual ACK lysis buffer, and centrifuged again at 500 x *g* for 5 min at 4°C before aspirating the supernatants.

The cells were resuspended into a single-cell suspension in MACS buffer and passed through a wetted LS column (Miltenyi Biotec) affixed to a QuadroMACS Magnet held by a QuadroMACS stand (both Miltenyi Biotec). The column was washed 3 times with 3 mL PBS each, and the initial flow-through and washes were collected to recover the non-magnetic cells. The LS column was then removed from the magnetic field, and the retained magnetic cell fraction was eluted with 1.5 mL PBS. The magnetic and non-magnetic cells were centrifuged at $500 \times g$ for 5 min at 4°C, and the supernatants were discarded. The cells were resuspended in PBS for cell counting and nanodiamond loading, as described above.

#### 1.5 Nanodiamond preparation and loading

Commercially available fluorescent nanodiamonds (Adámas Nanotechnologies) of 70 nm average size containing $\sim$ 3 ppm NV centers were used. Prior to incubation, nanodiamonds were dispersed by bath sonication for 10 min to minimize aggregation. Cells were incubated with nanodiamonds at a concentration of 40 $\mu$g/mL in PBS for 30 min at 37°C. Following incubation, cells were centrifuged at $500 \times g$ for 5 min at 4°C. The supernatant was discarded, and cells were washed twice with 200 $\mu$L PBS to remove excess extracellular nanodiamonds. The final cell pellet was resuspended in 100 $\mu$L of MACS buffer (PBS supplemented with 0.5% bovine serum albumin and 2 mM EDTA) for imaging.

#### 1.6 Quantum sensing measurements

Cell suspensions were mounted in imaging chambers constructed by sealing 80 $\mu$L of cells between two cleaned glass coverslips with vacuum grease or epoxy. Measurements were conducted at room temperature using a home-built confocal microscope equipped with a 520 nm continuous-wave laser for NV center excitation. A galvo mirror scanning system was used to perform fluorescence raster imaging to localize nanodiamond clusters inside cells. ODMR spectra were recorded by sweeping the frequency of the applied microwave field while monitoring fluorescence intensity. $T_1$ spin-lattice relaxation measurements were performed using a standard inversion-recovery pulse sequence. For each cell, 5–6 measurements were performed and averaged to improve signal reliability. Brightfield images were obtained using a white light LED source integrated into the same optical path to confirm cell integrity and intracellular localization of nanodiamond probes.

#### 1.7 Electronic and Spin Properties of NV Centers

Nitrogen-vacancy (NV) centers in diamond are point defects comprising a substitutional nitrogen atom adjacent to a lattice vacancy, replacing two carbon atoms in the diamond crystal lattice (*2*). This defect imparts the NV center with exceptional quantum spin properties, rendering it a highly versatile platform for nanoscale sensing.

The NV electronic ground state forms a spin-1 system with three spin sublevels: $|m_s = 0\rangle$ and $|m_s = \pm 1\rangle$. In the absence of external magnetic or electric fields, the $|m_s = \pm 1\rangle$ states are degenerate and separated from the $|m_s = 0\rangle$ state by a zero-field splitting of 2.87 GHz (*3*). This energy separation enables optically detected magnetic resonance (ODMR), wherein sweeping an applied microwave frequency in the presence of laser excitation reveals spin transition dips in the NV fluorescence spectrum.

Spin initialization and readout are achieved optically using 520 nm green laser excitation. NV centers preferentially relax from the excited state into the $|m_s = 0\rangle$ ground state via intersystem crossing, resulting in spin-dependent fluorescence. The $|m_s = 0\rangle$ state produces higher fluorescence than the $|m_s = \pm 1\rangle$ states, allowing robust spin readout and initialization with high fidelity (~80–90%) (*4, 5*).

#### 1.8 Temperature Sensing with NV Centers

NV centers are exquisitely sensitive to environmental changes due to their strong spin-lattice coupling. One such application is nanoscale thermometry, enabled by the temperature dependence of the NV zero-field splitting parameter $D$.

As temperature increases, thermal expansion of the diamond lattice perturbs spin-spin interactions, producing a linear decrease in $D$, and hence a shift in the ODMR resonance frequency (*4, 5*). This linear behavior over physiological temperature ranges enables NV centers to act as precise intracellular thermometers.

Experimental demonstrations have achieved sub-kelvin resolution using ODMR-based NV thermometry, offering a powerful tool to probe thermal gradients and metabolic activity in living cells with nanoscale spatial resolution (*6, 7*).

#### 1.9 Magnetic Noise Sensing with NV Centers

In addition to temperature, NV centers are highly sensitive to local magnetic field fluctuations. The longitudinal spin relaxation time, $T_1$, of the NV electronic spin is influenced by nearby fluctuating magnetic fields, enabling detection of paramagnetic species and magnetic noise at the nanoscale.

This technique, known as NV relaxometry, relies on dipolar coupling between the NV spin and surrounding magnetic dipoles (e.g., unpaired electrons in $Fe^{3+}$, $Mn^{2+}$, or free radicals), which induces spin-lattice relaxation and shortens $T_1$ (*8, 9*). The resulting change in fluorescence contrast during a pulsed relaxation measurement serves as a proxy for the local magnetic environment.

Recent advances have enabled detection of nanomolar concentrations of paramagnetic species in liquid environments using NV-based relaxometry. In our recent work (*10*), we utilized optically trapped fluorescent nanodiamonds (FNDs) to stabilize NV positions in solution, improving measurement reproducibility and mitigating artifacts from Brownian motion and charge-state fluctuations. This approach enhances the reliability of NV relaxometry in biological contexts.

#### 1.10 Temperature Calibration Using ODMR Center Frequency

The zero-field splitting (ZFS) parameter of nitrogen-vacancy (NV) centers in diamond is sensitive to temperature due to thermal expansion of the lattice, which perturbs the spin–spin interactions within the NV electronic ground state (*5*). As temperature increases, the ZFS decreases linearly, resulting in a measurable shift of the ODMR center frequency. This property provides the basis for NV-based quantum thermometry in physiological environments.

To calibrate the relationship between temperature and ODMR center frequency in our system, we conducted a controlled heating experiment. Fluorescent nanodiamonds (FNDs) containing NV centers (average diameter: 70 nm) were dispersed on a glass coverslip mounted over a printed circuit board (PCB) equipped with a resistive microheater. Temperature was varied from room temperature ( 25°C) to 38°C in  1–2°C increments. For each temperature point, fluorescence spectra were collected from five spatially distinct individual FNDs.

Each ODMR spectrum was fit to a Two-term Lorentzian fitting to extract the central resonance frequency. The five center frequencies obtained at each temperature were averaged to yield the temperature-dependent ODMR shift, and the standard deviation was used to calculate the error. The final calibration curve is plotted in Fig. S3.

We observed a linear dependence of center frequency on temperature, with a slope of −68.2 ± 8.0 kHz/°C and an intercept of 2.871 ± 0.0002 GHz. This linear model was used throughout the main manuscript to convert measured ODMR center frequencies into intracellular temperatures.

Importantly, we note that we verified that temperature measurements of our nanodiamond clusters in PBS showed that under our imaging conditions, the fitted temperature obtained from the measured ODMR spectrum 25.6 ± 3.8 °C (N = 8) was consistent with the thermocouple-measured room temperature 24.6 °C. Furthermore, in a separate control test ODMR measurements taken before and after hours-scale continuous laser illumination of nanodiamond clusters in magnetic splenocytes treated with FCCP resulted in a fitted center frequency drift rate of 0.01 kHz/s (N = 3), which would correspond to an apparent temperature drift of 0.1 °C under our ODMR protocol (*11, 12*).

#### 1.11 NV Relaxometry Protocol for Magnetic Noise Sensing

To probe the magnetic environment of intracellular NV centers, we implemented a differential $T_1$ relaxometry sequence adapted for nanodiamond quantum sensing (Fig. S4). This protocol enables robust extraction of the longitudinal spin relaxation time ($T_1$), which is sensitive to local magnetic noise arising from paramagnetic species.

The sequence consists of four key stages: (I) a 100 µs dark interval to establish charge state equilibrium, (II) a 5 µs green laser pulse ($\lambda$ = 520 nm) for spin polarization into the $|m_s = 0\rangle$ state, (III) a variable relaxation time interval $t$, and (IV) a readout laser pulse for fluorescence detection. Fluorescence is collected in a 1 µs detection window and recorded as the bright signal $S_B(t)$. To cancel out signal offsets and improve sensitivity to spin population differences, a second acquisition is performed using an identical sequence but with a $\pi$-pulse applied immediately before the final readout. This generates a corresponding dark signal $S_D(t)$.

The differential contrast is computed as:

$$C(t) = \frac{S_B(t) - S_D(t)}{(S_{Ref1}(t) + S_{Ref2}(t))/2},$$

where $S_{Ref1}(t)$ and $S_{Ref2}(t)$ are reference fluorescence signals collected at the end of the initialization laser pulse.

This differential protocol mitigates photoionization artifacts and laser-intensity dependencies, which are known to confound all-optical relaxometry measurements in nanodiamond systems (*13–15*). In dense NV ensembles or cellular environments, the resulting decay curves often follow a stretched exponential form, reflecting a distribution of local magnetic environments within individual FNDs (*16*).

This method enables robust, quantitative readout of magnetic noise levels with high reproducibility, forming the basis for magnetic sensing throughout our study.

### Figures S1 to S4

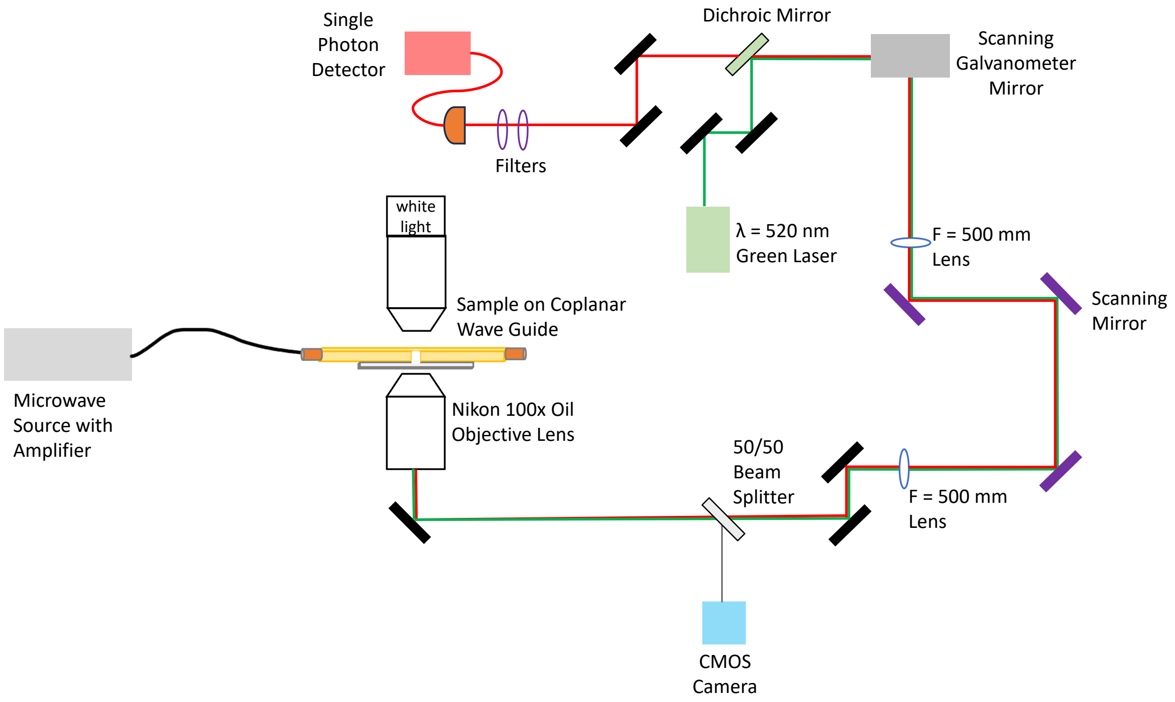


**Figure S1**: Schematic of optical setup.

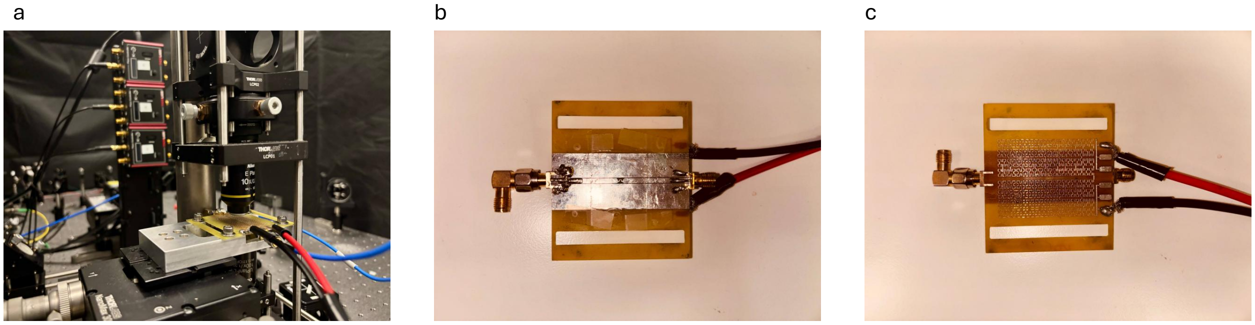


**Figure S2**: **Integrated Microwave and Thermal Control Platform for NV Quantum Sensing.** (a) Cells with internalized nanodiamonds are sealed on a coverglass and mounted onto a printed circuit board (PCB) featuring a coplanar waveguide (CPW) for microwave delivery. (b) The CPW contains an Ω-shaped loop (radius 150 μm) with a central through-hole for white-light imaging. (c) A patterned resistive heater on the back side of the PCB enables temperature control during measurements.

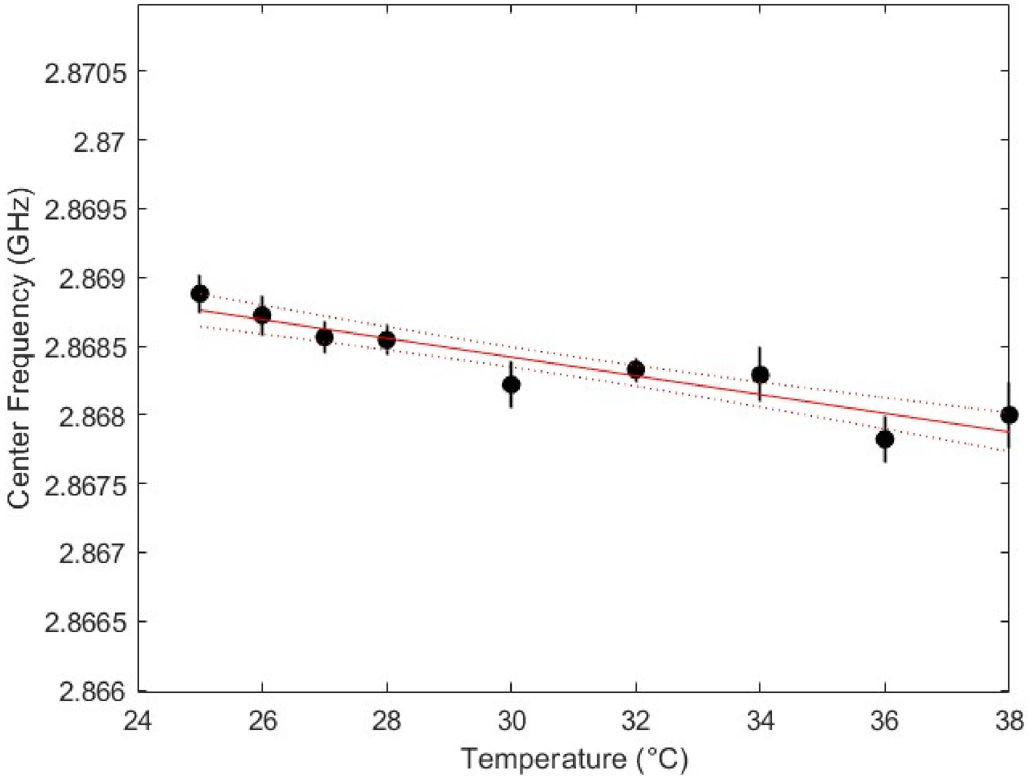


**Figure S3**: **Calibration of NV center frequency as a function of temperature.** ODMR center frequency was measured for five independent nanodiamonds (NDs) at each temperature point and averaged. A linear fit (solid red line) yielded a temperature sensitivity of −68.2 ± 8.0 kHz/°C. Dotted lines denote the 95% confidence interval of the fit.

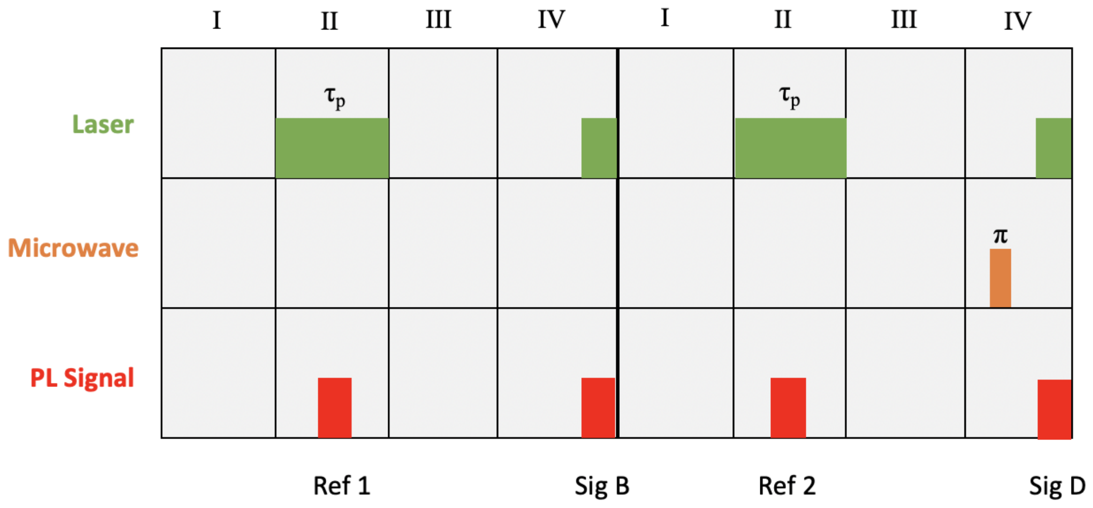


**Figure S4**: **Pulse sequence for NV relaxometry.** (I) Dark interval to establish charge state equilibrium. (II) Spin polarization with a 5 µs 520 nm laser pulse. (III) Free relaxation interval $t$. (IV) Readout window (1 µs) to collect fluorescence. Each trace is measured both with and without a final $\pi$-pulse. Contrast is computed from the differential signal to extract $T_1$ spin relaxation dynamics.
